# ADP-ribose triggers neuronal ferroptosis by rewiring purine and pyrimidine metabolism

**DOI:** 10.1101/2021.05.12.443941

**Authors:** Haoqi Ni, Yingying He, Peng Cui, Hebing Chen, Guoqing Lv, Huiyan Lei, Feng Wang, Qichuan ZhuGe, Baodong Chen, Ling Liang, Yong Zhang, Fuping You, Lin Yuan

**Author notes:** These authors contributed equally to this work (Haoqi Ni, Yingying He, Peng Cui, Hebing Chen, Lin Yuan). Corresponding author. Running title: ADP-ribose induces neuronal ferroptosis.

## Abstract

Hyperactivation of NAD^+^-consuming pathways frequently occurs in neurological diseases, however therapies replenishing NAD^+^ levels show limited therapeutic efficacy, indicating more complex underlying pathophysiology. Here, we delineate a pathogenic link between ADP-ribose —a product of NAD^+^ consumption—and a metabolic rewiring-dependent form of neuronal ferroptosis. We demonstrate that oxidative stress induces neurons to produce ADP-ribose through the PARP1-PARG axis. ADP-ribose directly binds and inhibits the equilibrative nucleoside transporter ENT2, remodeling *de novo* purine and pyrimidine synthesis by hyperactivating the inosine-hypoxanthine-xanthine oxidase and glutamine-dihydroorotate-dihydroorotate dehydrogenase axes. This overproduces superoxide radicals and drives lipid peroxidation and neuronal ferroptosis. Elevated ADP-ribose levels were observed in neurological disease models, and acute ADP-ribose exposure severely reduced mouse brain neurons *in vivo*. Critically, interventions blocking ADP-ribose signaling alleviated cognitive decline in mouse intracerebral hemorrhage models. Our findings characterize ADP-ribose signaling as linking NAD^+^ consumption to neuronal ferroptosis, and provide a theraputic strategy for neuropathologies involving NAD^+^ consumption and oxidative stress.

## INTRODUCTION

Maintenance of nicotinamide adenine dinucleotide (NAD^+^) homeostasis through its intracellular physiological level between 0.2 and 0.5 mM is essential for neuronal redox and bioenergetic homeostasis ^1^. However, hyperactivation of the major NAD^+^-consuming enzyme families— poly(ADP-ribose) polymerases (PARPs), Sirtuins, and NAD nucleosidases (*e.g.*, CD38, CD157, and SARM1)—is associated with several neurological disorders, including Alzheimer’s disease (AD) and hemorrhagic stroke ^1,2^. These NAD^+^-consuming enzyme families ultimately consume NAD^+^ to yield ADP-ribose and release nicotinamide ^3^. While supplementation of NAD^+^ or its precursors shows limited therapeutic benefit in patients with hyperactivated NAD^+^-consuming enzymes ^3^, indicating more complex underlying pathological mechanism.

Under oxidative stress conditions, increases in activity of PARP1 lead to upregulation of NAD^+^ consumption and protein poly-ADP-ribosylation ^4^. Poly-ADP-ribosylation is reversible and can be degraded by poly(ADP-ribose) glycohydrolase (PARG) into ADP-ribose polymers (“PAR”) and ADP-ribose monomers (“ADP-ribose”) ^5^. PAR can induce mitochondrial apoptosis-inducing factor (AIF) release and nuclear translocation, leading to “parthanatos” cell death ^6^. However, PAR is rapidly degraded by PARG into ADP-ribose ^5^. The function of ADP-ribose remains unclear, although some studies show PARG knockout mice exhibit higher brain damage after stroke^7^, whereas others report PARG inhibitors can rescue neuronal cell death induced by the glutamate receptor agonist NMDA^8^. Collectively, these findings suggest that both PAR and ADP-ribose accumulation may influence neuronal cell death.

Ferroptosis is a regulated form of cell death that depends on iron and Lands’ cycle, characterized by excessive lipid peroxidation^9^. Lipid peroxidation is initiated by the formation of a carbon-centered radical on polyunsaturated fatty acyl phospholipids (PUFA-PLs), which can be seed by hydroxyl or peroxyl radicals generated from iron-catalyzed Fenton reactions ^10^. If not converted to a lipid hydroperoxide and subsequently reduced to the corresponding alcohol, the propagation of free-radical-mediated reactions will commence, ultimately resulting in a myriad of secondary products, disruption of membrane integrity, and membrane rupture with the release of cellular contents ^11^. Due to their high PUFA-PL content, neuronal menbranes are particularly susceptible to peroxidation ^11^. While cells have evolved multiple defenses against ferroptosis by increasing antioxidant capacity, mainly via the glutathione peroxidase 4 (GPX4)/glutathione (GSH) system^12–14^, the ferroptosis suppressor protein 1 (FSP1)/coenzyme Q (CoQ) system^15,16^, and the GTP cyclohydrolase-1 (GCH1)/tetrahydrobiopterin system^17^, intracellular metabolic dysregulation causing redox imbalance appears to be the essential trigger of lipid hydroperoxide and ferroptosis.

Here, we found that exposure to ferroptosis inducers promoted the accumulation and release of ADP-ribose from cultured primary neurons. Using thermal proteome analysis (TPP) and mass spectrometry-based metabolomics, we show that extracellular ADP-ribose directly binds to the equilibrative nucleoside transporter 2 (ENT2) protein, inhibiting its bidirectional transport of purine and pyrimidine nucleosides. This transport inhibition led to overactivation of the inosine-hypoxanthine-xanthine oxidase (XO) and glutamine-dihydroorotate-dihydroorotate dehydrogenase (DHODH) axes in neurons, resulting in excessive superoxide anion production and lipid peroxidation-dependent ferroptotic death of neurons. Microinjecting ADP-ribose into mouse brains dramatically reduced neuron numbers, an effect rescued by co-administering XO/DHODH inhibitors. Furthermore, intracerebral hemorrhage (ICH) induced production in mouse neurons *in vivo* with subsequent cognitive decline, aspects mitigated by attenuating ADP-ribose production via PARP1 knockdown or microinjecting the ADP-ribose pyrophosphatase Nudix hydrolase 9 (NUDT9).

## RESULTS

### Ferroptotic metabolite ADP-ribose induces neuronal cell death and is elevated in neurological disease models

Given that apoptotic cells release metabolites that can impact neighboring tissues ^18,19^, we investigated whether metabolites released from ferroptotic neurons affect unstressed neurons. Primary cortical neurons isolated from E16.5 mice (Fig. S1A) were cultured *in vitro* for 2 weeks to mature and then exposed for 6 hours to 100 µM hemin^20,21^, a ferroptosis inducer that model intracerebral hemorrhage *in vitro* (Fig. 1A). The medium was subsequently replaced with fresh medium and incubated for an additional 6 hours, at which point a CCK-8 assay showed ∼70% cell death (Fig. S1B). When this conditioned medium was collected and concentrated ten-fold, despite the absence of detectable hemin, it induced cell death upon addition to mature primary cortical neuron cultures (Fig. 1B). This observation suggests ferroptotic neurons release extracellular factor(s) inducing death in unstressed neurons. To determine if the released factor(s) was proteinaceous, we heat-treated the conditioned medium at 95°C for 5 minutes before adding to mature primary cortical neuron cultures, which did not prevent cell death (Fig. 1, A and B). Fractionation by molecular sieve indicated the factor(s) were <1 kDa (Fig. 1, A and B). Untargeted metabolomics identified ADP-ribose as the small molecule most upregulated (> 90-fold) in the medium after hemin exposure (Fig. S1C and Supplementary Table 1). MS-based quantification using an authentic standard curve confirmed significantly higher intracellular and extracellular ADP-ribose in hemin-treated neurons (Fig. 1, C and D and Fig. S1D).

**Fig. 1.**
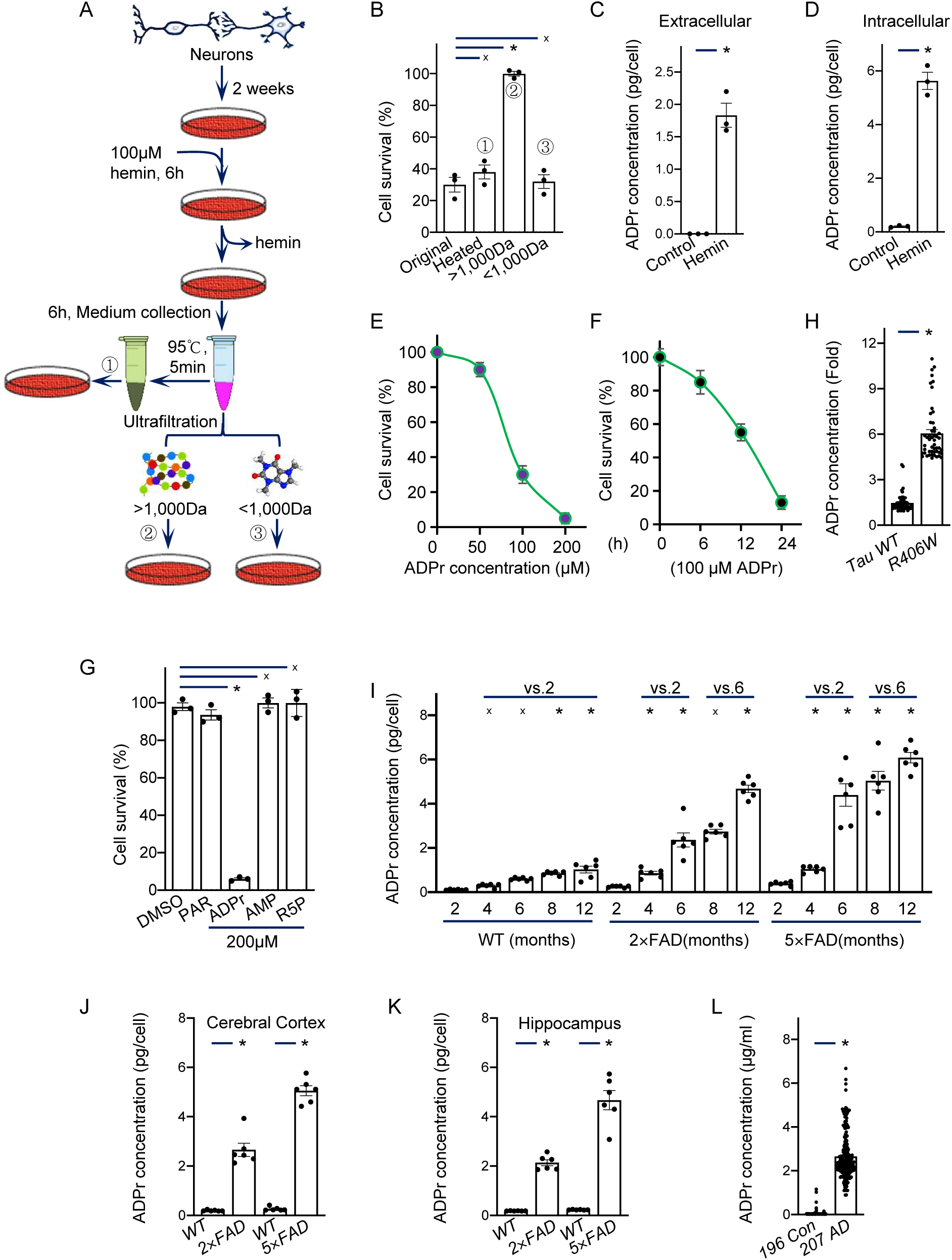
ADP-ribose is toxic to neurons and upregulated in neurological diseases A. Schematic conditioned medium preparation. EMX-A14 murine primary cortical neurons were treated with 100 μM hemin for 6 hours, followed by replacement with fresh medium for 6 hours. Conditioned medium was concentrated (original), heated or filtered to generate fractions (>1,000 Da or <1,000 Da). B. Primary cortical neurons were treated with conditioned medium for 24 hours and cell viability assessed by CCK-8 assay. Kruskal–Wallis one-way ANOVA followed by Dunn’s multiple comparison tests. C. Mature primary cortical neurons treated with 100 μM hemin for 6 hours. Extracellular ADP-ribose levels determined by LC-MS/MS (QTRAP 6500). Student’s t-test with Welch’s correction. D. Intracellular ADP-ribose levels in mature primary cortical neurons treated with 100 μM hemin for 6 hours. Student’s t-test with Welch’s correction. E. Viability of primary cortical neurons (plating 24 hours) was assessed after 24 hours treatment with ADP-ribose doses. F. Viability of mature cortical neurons (plating 2 weeks) treated with 100μM ADP-ribose over time. G. Viability of mature primary cortical neurons exposed to PAR, ADP-ribose, AMP, or ribose 5-phosphate (R5P) for 24 hours. Kruskal–Wallis one-way ANOVA followed by Dunn’s multiple comparison tests. H. Relative ADP-ribose levels in tauR406W and wild-type fruit fly brains measured by QTRAP 6500 LC-MS/MS. n=50 flies. Mann–Whitney test. I. Hippocampal neuronal ADP-ribose levels in APP/PS1 and 5×FAD mice vs age. n=6. Two-way ANOVA (mouse strain × age) followed by Bonferroni posthoctests. J. Cortical neuronal ADP-ribose levels in 6-month-old APP/PS1 and 5×FAD mice and wild-type littermate controls. n=6. Kruskal–Wallis one-way ANOVA followed by Dunn’s multiple comparison tests. K. Hippocampal neuronal ADP-ribose levels in 6-month-old APP/PS1 and 5×FAD mice and wild-type littermate controls. n=6. Kruskal–Wallis one-way ANOVA followed by Dunn’s multiple comparison tests. L. Cerebrospinal fluid (CSF) ADP-ribose levels were determined by QTRAP 6500 LC-MS/MS in healthy controls (n= 196) and patients with AD (n= 207). Mann–Whitney test. All data are means ± SEM. *, P< 0.01; x, no significant difference. See also Figures S1 and S2. Statistics source data can be found in Supplementary Table 2.

We next tested if ADP-ribose directly induce neuronal cell death. ADP-ribose exposure killed murine naïve primary cortical neurons (24h post-plating), mature primary cortical neurons (2 weeks post-plating), murine hippocampal HT22 neurons, and human SH-SY5Y neuroblastoma cells (Fig. 1, E and F and Fig. S1, E and F), with slight differences in sensitivity. In contrast, ADP-ribose exposure did not kill murine primary astrocytes (Fig. S1G). Furthermore, neither ADP-ribose precursors (PAR) nor its degradation products (AMP and R5P) induced neuronal death (Fig. 1G and S1H). Moreover, cell death was not observed when the conditioned medium was treated with ADP-ribose pyrophosphatase NUDT9 prior to assays, corroborating ADP-ribose specificity (Fig. S2, A to D). Additionally, ADP-ribose sensitized primary cortical neurons to hemin- and L-homocysteic acid-induced cell death (Fig. S2E). Collectively, these data indicate ADP-ribose exposure triggers neuronal cell death.

We subsequently explored the potential *in vivo* relevance of stress-induced ADP-ribose production in neurons in multiple neurological disease models. LC-MS/MS quantification of brains from the well-characterized TauR406W *Drosophila melanogaster* model, which displays early-onset familial dementia features ^22^, showed elevated ADP-ribose levels versus wild-type flies (Fig. 1H). Furthermore, we examined brain cells from Alzheimer’s disease (AD) mouse models, specifically APP/PS1 and 5×FAD mice ^23,24^. While hippocampal neuronal ADP-ribose gradually increased with age in wild-type mice, it increased substantially more in APP/PS1 and 5×FAD mice (Fig. 1I). Compared to age-matched wild-type littermates, both transgenic lines exhibited significantly higher ADP-ribose levels in cerebral cortical and hippocampal neurons (Fig. 1, J and K). Additionally, cerebrospinal fluid (CSF) samples from AD and control subjects showed elevated median ADP-ribose in AD patients (n=207, 2.53 μg/ml) versus controls (n=196, 0.03 μg/ml, *P*<0.0001) (Fig. 1L). Hence, the presence of elevated brain ADP-ribose in neurological disease models implicates the potential pathological relevance of ADP-ribose.

### ADP-ribose production is induced by the PARP1-PARG axis

We investigated how ADP-ribose is generated in ferroptotic neurons. Exposure of neurons to hemin reduced GSH levels and induced H_2_O_2_ generation (Fig. S3A). Under oxidative stress, PARP1 is activated and utilizes NAD^+^ as a substrate to catalyze protein poly-ADP-ribosylation, which is rapidly degraded by PARG to yield ADP-ribose, contributing to the intracellular ADP-ribose pool ^8,25–27^. We found that primary cortical neurons from *Parp1* neuron-specific knockout mice exhibited reduced NAD^+^ consumption and ADP-ribose production upon hemin exposure compared to wild-type controls (Fig. 2, A and B; fig. S3B). Moreover, under cysteine withdrawal, ADP-ribose production was significantly reduced in PARP1 knockout neurons compared to wild-type (Fig. 2C). To confirm PARP1’s role in production of ADP-ribose precursors, we pretreated neurons with the PARG inhibitor PDD to block rapid poly-ADP-ribosylation hydrolysis. We observed that exposure to hemin, erastin and glutamate significantly increased poly-ADP-ribosylation (Fig. 2D and fig. S3C), which was reduced by the radical scavenger α-TOC (Fig. S3C). Additionally, the PARP1 degrader iRucaparib-AP6^28^ diminished hemin-, erastin- and glutamate-induced poly-ADP-ribosylation (Fig. 2D; fig. S3, D to F). These results indicate that activated PARP1 promotes ADP-ribose production under ferroptosis-induced oxidative stress.

**Fig. 2.**
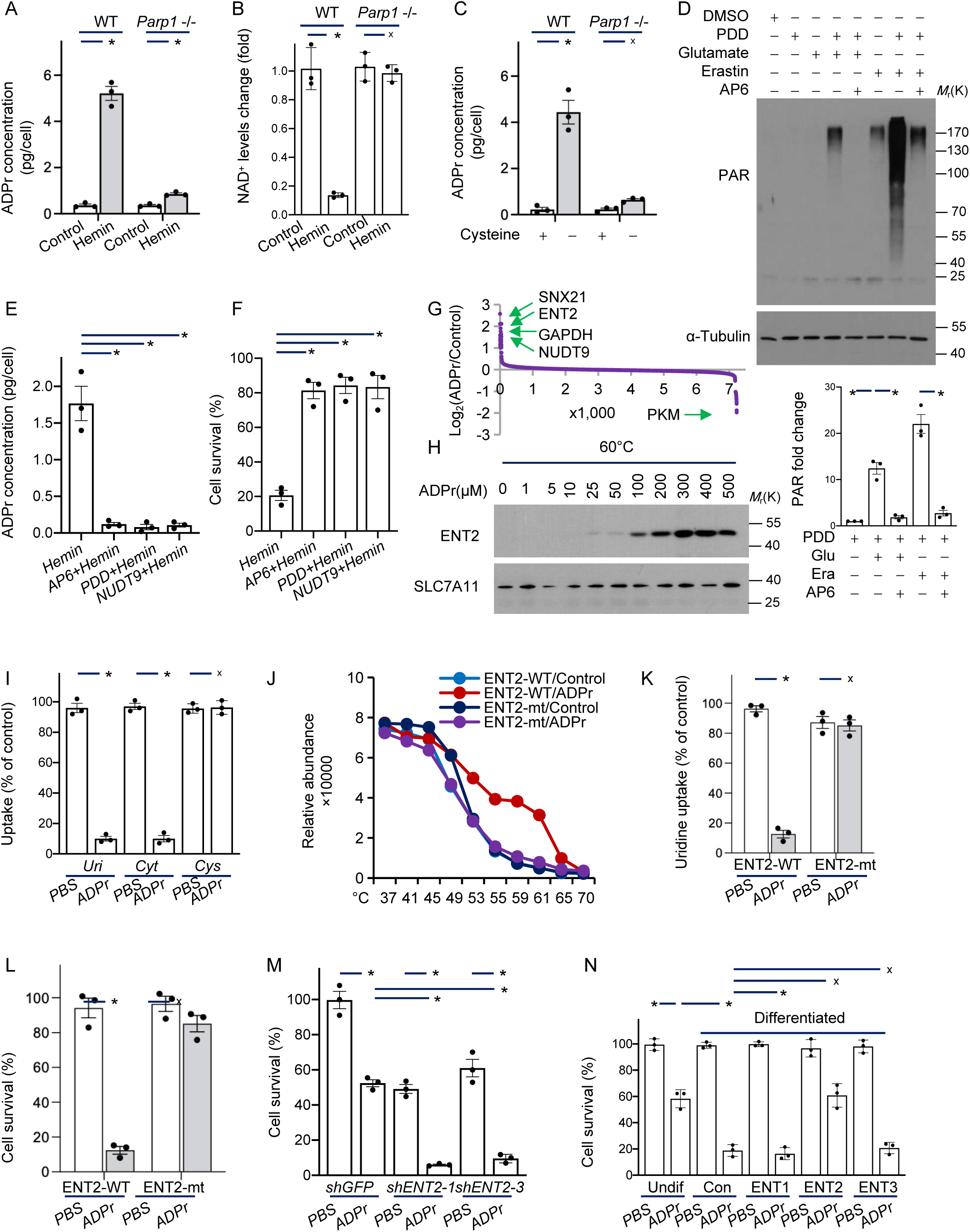
**ADP-ribose is produced by the PARP1-PARG axis and directly targets ENT2** A. Primary cortical neurons isolated from the mice with neuron-specific *Parp1* knockdown were treated with 100 μM hemin. After 6 hours, the intracellular ADP-ribose levels were determined by QTRAP 6500 LC-MS/MS. B. Primary cortical neurons isolated from the mice with neuron-specific *Parp1* knockdown were treated with 100 μM hemin. After 6 hours, the NAD^+^ levels were measured by NAD/NADH Assay Kit (ab65348). C. Primary cortical neurons isolated from the mice with neuron-specific *Parp1* knockdown were cultured for 12 hours in cystine-free media. The intracellular ADP-ribose levels were determined by QTRAP 6500 LC-MS/MS. D. Hippocampal HT22 neurons were pretreated with 0.1μM iRucaparib-AP6 (PROTAC PARP1 degrader) or 2μM PDD (PARG inhibitor) for 1 hour, then treated with 5μM erastin or 5mM glutamate. After 8 hours, the cell lysates were subject to western blotting with anti-PAR antibodies. Quantification of total PAR levels in neuronal cells under oxidative stress (below). E. Primary cortical neurons were pre-treated with 0.1μM iRucaparib-AP6 or 2μM PDD for 1 hour, then exposed to 100μM hemin for 6 hours, followed by culture in hemin-free medium for 6 hours. Conditioned medium from lane 4 was treated with 1 µg/ml ADP-ribose hydrolase NUDT9 (ab107153) to eliminate ADP-ribose prior to measurement of its concentration. F. Primary cortical neuronal viability assessed after culture with conditioned media from (E) for 24 hours. G. Total SH-SY5Y proteins treated with or without 10μM ADP-ribose and analyzed by TMT10 for thermal stability changes. H. Recombinant ENT2 or SLC7A11 incubated with ADP-ribose doses for 10 mins at room temperature then denatured at 60°C for 3 min and western blotted for stability. I. ^13^C-uridine (Uri, 50 μM) or ^13^C-cytidine (Cyt, 50 μM) uptake by SH-SY5Y cells pretreated with 100 μM ADP-ribose for 3 hours. ^13^C-cystine (Cys, 50 μM) as a negative control. J. Detection of binding between *E. coli*-derived ENT2 wild-type (WT) or Q76A/Q145A mutant (mt) and ADP-ribose by thermal shift assay. K. HT22 cells stably expressing ENT2-WT or ENT2-mt were pretreated with 100 μM ADP-ribose for 3 hours, and the uptake of ^13^C-uridine (50 μM) was measured. L. HT22 cells stably expressing ENT2-WT or ENT2-mt were treated with 100 μM ADP-ribose for 24 hours and cell viability was measured by CCK-8 assay. M. SH-SY5Y cells stably expressing the indicated shRNAs were treated with 100μM ADP-ribose for 24 hours and cell viability assessed by by CCK-8 assay. N. SH-SY5Y cells were differentiated through 6 days of treatment with 10 μM all-trans-retinoic acid (RA). Differentiated SH-SY5Y cells expressing the indicated plasmids were treated with 100μM ADP-ribose for 24 hours before assessing cell viability. Data are means ± SEM of three independent experiments. Kruskal–Wallis one-way ANOVA followed by Dunn’s multiple comparison tests. *, P< 0.01; x, no significant difference. See also Figures S3 to S5. Statistics source data can be found in Supplementary Table 2.

We further tested whether inhibition of the PARP1-PARG axis could prevent hemin-induced ADP-ribose generation. Specifically, murine primary cortical neurons were pre-treated with PDD or iRucaparib-AP6 for 1 hour followed by 6 hours of exposure to 100 µM hemin and a further 6 hours of culture in hemin-free medium (Fig. S3G). Although significant cell death was still detected by CCK-8 assay (Fig. S3H), no ADP-ribose was detected in the conditioned medium and no additional cell death was observed upon treating murine primary neuron cultures with this conditioned medium (Fig. 2, E and F). These results suggest that ferroptosis-stressed neurons generate ADP-ribose through the PARP1-PARG axis, with subsequent ADP-ribose release due to membrane integrity disruption.

### ADP-ribose targets and inhibits the nucleoside transporter ENT2

To uncover potential intracellular targets of ADP-ribose, we employed thermal proteome profiling (TPP), which can unbiasedly identify ligand-protein interactions based on increased protein thermal stability upon complex formation^29^. Notably, intracellular ADP-ribose levels did not increase in SH-SY5Y cells after exposure to semi-lethal doses (Fig. S3I), suggesting ADP-ribose binds plasma membrane proteins to trigger cell death. Therefore, we treated total SH-SY5Y proteins with ADP-ribose and performed TPP, which identified a total of 7,263 proteins. Among the top 5-ranking candidates with >2-fold thermal stability increases upon ADP-ribose treatment, the equilibrative nucleoside transporter 2 (ENT2), also known as SLC29A2, stood out prominently as a plasma membrane protein (Fig. 2G and Supplementary Table 2). Furthermore, we validated these results by treating murine primary neurons with varying doses of ADP-ribose and detecting a dose-dependent increase in ENT2 thermal stabilization (Fig. 2H). In addition, direct ADP-ribose binding to *E. coli*-purified ENT2 was observed via microscale thermophoresis and thermal melting assays (Fig. S4, A and B). However, no ENT2 binding was detected to either PAR or di-ADP-ribose in thermal melting assays, implicating ADP-ribose specifically (Fig. S4C).

ENT bidirectionally transports various purine and pyrimidine nucleosides, including uridine, cytidine, inosine and hypoxanthine ^30–32^. Using the ENT1 crystal structure (PDB:6OB6)^33^ and an AlphaFold2 model, we performed *in silico* docking suggesting that ADP-ribose occupies the ENT2 nucleoside transport pocket (Fig. S4, D to F). ^13^C-metabolic flux analysis demonstrated that ADP-ribose pretreatment abolished uptake of substrates like uridine and inosine into SH-SY5Y cells without inducing cell death (Fig. 2I, Fig. S4G and Supplementary Table 3). High-dose nitrobenzylthioinosine (NBMPR, 100 μM), an ENTs inhibitor, also inhibited substrate uptake (Fig. S4H). Export of substrates from SH-SY5Y cells was also abolished upon ADP-ribose pretreatment (Fig. S4, I and J). Metabolomics revealed that ADP-ribose exposure reduced intracellular uridine and cytidine, while increasing inosine and hypoxanthine levels (Fig. S4, K and L). Thus, extracellular ADP-ribose inhibits ENT2-mediated transport of pyrimidine and purine nucleosides.

Mutations in ENT2 residues Q76/Q145 attenuated ADP-ribose binding *in vitro* (Fig. 2J), and abolished substrate transport impairment by ADP-ribose in HT22 cells (Fig. 2K, and Fig. S5A). HT22 cells expressing mutant ENT2 were resistant to ADP-ribose-induced death versus wild-type cells (Fig. 2L). The ENT2 inhibitors dipyridamole and high-dose NBMPR partially reduced primary neuron viability, implicating ENT2 inhibition induces neuronal death (Fig. S5B). ENT2, but not ENT1, knockdown sensitized SH-SY5Y cells to ADP-ribose-induced cell death (Fig. 2M and Fig. S5, C to G). Retinoic acid (RA)-differentiated SH-SY5Y cells^34^, which downregulate ENT2 (Fig. S5, H and I), exhibited heightened sensitivity to ADP-ribose-induced death versus undifferentiated cells (Fig. 2N). Stably expressing ENT2, but not ENT1/3, rendered differentiated cell resistance to ADP-ribose (Fig. 2N, and Fig. S5J). Collectively, these results indicate that ADP-ribose triggers neuronal cell death by inhibiting ENT2.

### ADP-ribose is a neuronal ferroptosis inducer

We further characterized the cell death pathway(s) induced by ADP-ribose in neurons using inhibitors of distinct cell death forms. ADP-ribose-mediated cell death was fully rescued by ferroptosis inhibitors including the iron chelator deferoxamine (DFO) and the radical-trapping antioxidants—α-Tocopherol (α-Toc), ferrostatin-1 (Fer-1) and liproxstatin-1 (Lip-1) (Fig. 3A and Fig. S6, A and B). Inhibitors of autophagy or apoptosis did not exert protective effects (Fig. 3A and Fig. S6, A and B). Additionally, ADP-ribose treatment increased levels of the lipid peroxidation marker 4-HNE in murine primary cortical neurons (Fig. 3B). Cell death induced by ENT2 knockdown or inhibition with NBMPR was also rescued by Fer-1 or Lip-1 (Fig. 3C and S6C). Together, these results demonstrate that ADP-ribose exposure triggers neuronal ferroptosis.

**Fig. 3.**
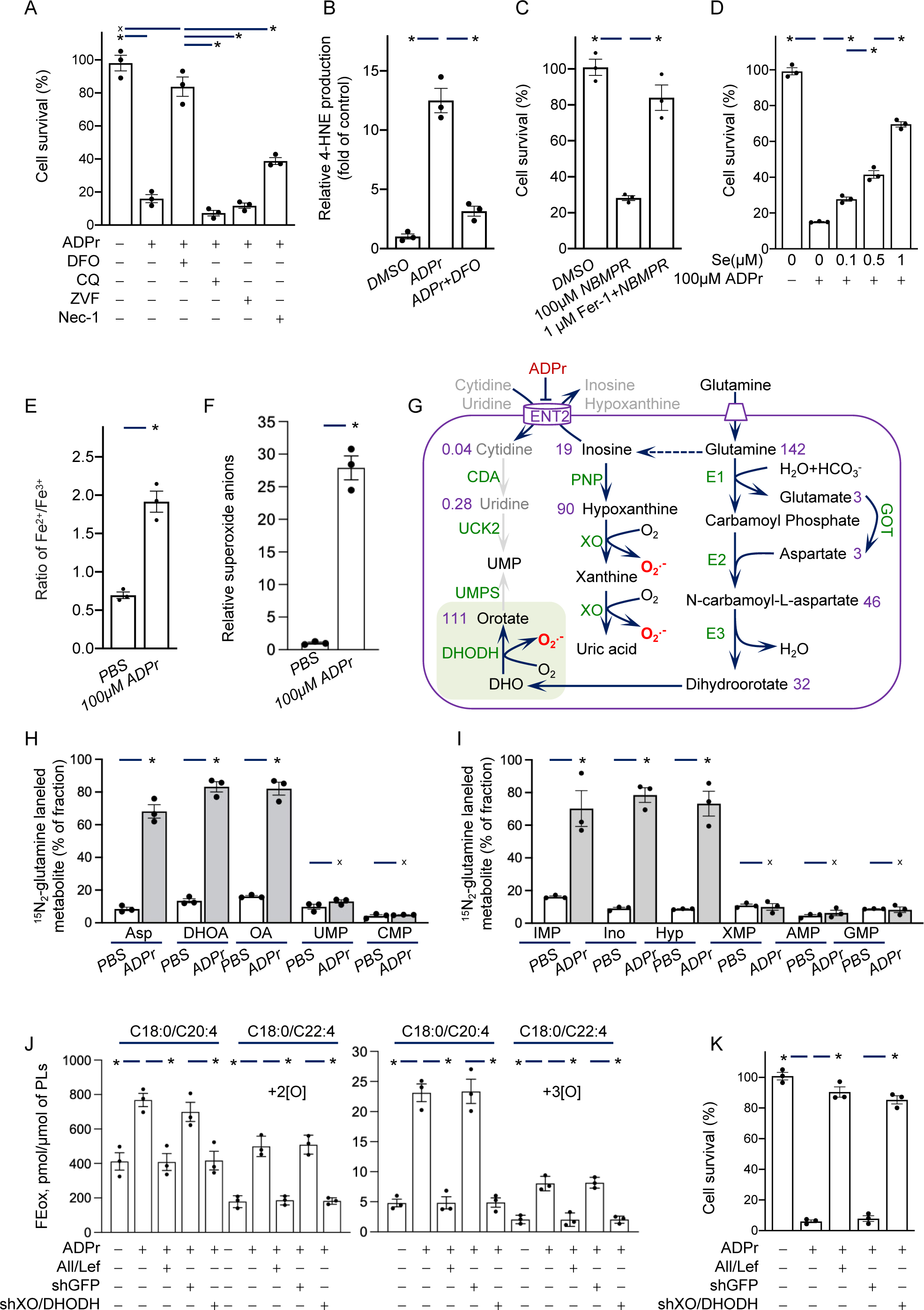
**ADP-ribose triggers neuronal ferroptosis in a XO/DHODH-dependent manner** A. Quantification of primary cortical neurons viability after 24 hours of indicated treatment. ADP-ribose, 100μM; DFO, deferoxamine, a ferroptosis inhibitor, 100μM; CQ, chloroquine, an autophagy inhibitor, 10μM; ZVF, Z-VAD-FMK, a pan-caspase inhibitor, 10μM; Nec-1, Necrostain-1, a necroptosis inhibitor, 10μM. B. 4-HNE levels in primary cortical neurons exposed to 100μM ADP-ribose for 12 hours, measured by 4-HNE assay kit (ab238538). C. Cultured mature cortical neuron viability at 24 hours with treatments. Fer-1, ferrostatin-1, a ferroptosis inhibitor. D. Cultured primary cortical neuron viability at 24 hours with treatments. E. Fe^2+^/Fe^3+^ ratio was measured in HT22 cells treated with 100μM ADP-ribose for 12 hours. F. Superoxide level was measured in HT22 cells treated with 100μM ADP-ribose for 12 hours. G. SH-SY5Y metabolomics identified altered purine and pyrimidine metabolites after 100μM ADP-ribose exposure for 12 hours. The change fold of metabolites under ADP-ribose treatment was labeled in purple number. H. Mass isotopomer analysis of aspartate (m+1), dihydroorotate (m+1/2), orotate (m+1/2), UMP (m+1/2) and CMP (m+1/2) in HT22 cells cultured with 3 mM ^15^N_2_-glutamine and 100μM ADP-ribose for 12 hours. I. Mass isotopomer analysis of IMP (m+2/3), inosine(m+2/3), hypoxanthine(m+2/3), XMP (m+2/3), AMP (m+2/3) and GMP (m+2/3/4) in HT22 cells cultured with 3 mM ^15^N_2_-glutamine and 100μM ADP-ribose for 12 hours. J. SH-SY5Y cells stably expressing the indicated shRNAs or pre-treated with 10μM allopurinol and leflunomide for 1 hour, were treated with 100μM ADP-ribose for 12 hours. The levels of di- and tri-oxygenated PE-(C18:0/C20:4) and PE-(C18:0/C22:4) were detected by MS/MS. K. Differentiated SH-SY5Y cells stably expressing the indicated shRNAs or pre-treated with 10μM allopurinol and leflunomide for 1 hour were treated with 100μM ADP-ribose and the cell viability was tested after 24 hours. Data are means ± SEM of three independent experiments. Kruskal–Wallis one-way ANOVA followed by Dunn’s multiple comparison tests in A-D and H-J. Student’s t-test with Welch’s correction in E and F. *, P < 0.01; x, no significant difference. See also Figures S6 to S9. Statistics source data can be found in Supplementary Table 2.

Notably, low-dose ADP-ribose treatment exacerbated cortical neuron death induced by sublethal doses of various ferroptotic inducers, including hemin (an oxidized form of heme), RSL3 (a GPX4 inhibitor), or FIN56 (an agent that activates squalene synthetase), which was rescued by Fer-1 (Fig. S6D). GPX4 converts lipid hydroperoxides to lipid alcohols and prevents ferroptosis ^13^, and its expression is dependent on selenium availability (Fig. S6, E and F) ^21^. Selenium dose-dependently abrogated ADP-ribose-induced neuronal death (Fig. 3D). In addition, HT22 cells expressing Q76A/Q145A mutant ENT2 showed reduced lipid peroxidation and cell death after ADP-ribose treatment compared to wild-type cells (Fig.2L, and Fig. S6G). Thus, ADP-ribose triggers neuronal ferroptosis by inducing lipid peroxidation.

The initiation and propagation of lipid peroxidation are dependent on Fe^2+^-catalyzed Fenton reactions ^35^. Specifically, we observed that ADP-ribose exposure increased the Fe^2+^/Fe^3+^ ratio in HT22 cells (Fig. 3E), indicating that the Fe^3+^ generated after the reaction is reduced back to Fe^2+^. Cellular superoxide anions are known to modulate iron redox state and thereby induce ferroptosis ^9,36^. Indeed, superoxide anion assays confirmed that ADP-ribose exposure increased superoxide anion production in HT22 cells (Fig. 3F). While electron transport chain (ETC) dysfunction is typically associated with increased superoxide anions, ADP-ribose exposure had no direct effect on ETC activity (Fig. S6H). These results suggest that ADP-ribose triggers neuronal ferroptosis by accumulating superoxide anions.

### ADP-ribose triggers ferroptosis in a XO/DHODH-dependent manner

To investigate the mechanisms underlying ADP-ribose-induced superoxide anion accumulation, we performed untargeted metabolomics profiling of SH-SY5Y cells (Fig. S7, A to C; Supplementary Table 4). Treatment with ADP-ribose enriched key intermediates in both purine and pyrimidine biosynthesis pathways (Fig. 3G and Supplementary Table 5). Specifically, ADP-ribose treatment elevated intracellula levels of deoxyinosine, inosine and hypoxanthine, suggesting engagement of the inosine-hypoxanthine-XO purine catabolic pathway (Fig. 3G and S4L). Hypoxanthine can be oxidated by xanthine oxidase (XO) to generate superoxide anions^37^. Accordingly, XO knockdown or inhibition with allopurinol^38^ markedly reduced ADP-ribose-induced superoxide production (Fig. S8, A and B). Moreover, blocking hypoxanthine production through inhibition of purine nucleoside phosphorylase with forodesine ^39^ also protected murine primary cortical neurons from ADP-ribose cytotoxicity (Fig. S8C).

Additionally, levels of glutamine, dihydroorotate and orotate accumulated under ADP-ribose exposure, implicating activation of the glutamine-dihydroorotate-DHODH axis (Fig. 3G). Dihydroorotate dehydrogenase (DHODH) catalyzes the oxidation of dihydroorotate to orotate via ubiquinone with concomitant superoxide generation ^40^. Accordingly, inhibiting DHODH with either leflunomide or BAY-2402234, or depleting DHODH via shRNAs knockdown, significantly reduced ADP-ribose-induced superoxide production and lipid peroxidation levels (Fig. S8, C to E). Blocking glutamine uptake through GPNA treatment elicited comparable effects (Fig. S8C).

Glutamine, the primary precursor for *de novo* purine and pyrimidine synthesis, was essential for ADP-ribose-induced ferroptosis (Fig. S8F). Metabolic flux analyses using ^15^N_2_-glutamine revealed that ADP-ribose treatment substantially leads to intracellular accumulation of key purine (IMP, inosine and hypoxanthine) and pyrimidine (aspartate, dihydroorotate and orotate) intermediates derived from glutamine (Fig. 3, H and I; Fig. S9, A and D). Notably, downstream purine and pyrimidine nucleotides incorporated less of the ^15^N-label compared to precursors (Fig. Fig. 3, H and I; Fig. S9, A and D). This ADP-ribose-induced purine and pyrimidine *de novo* pathway can be explained by the reduced salvage pathway independently of metabolic enzyme regulation upon ADP-ribose exposure in ^13^C-flux analysis (Fig. S9, A to H). Collectively, these results demonstrate that ADP-ribose triggers ferroptosis by co-opting the XO and DHODH arms of purine and pyrimidine biosynthesis to generate cytotoxic ROS.

Simultaneous knockdown of XO and DHODH (Fig. S8A) or combined inhibition of XO/DHODH with allopurinol/leflunomide markedly reduced ADP-ribose-induced di- and tri-oxygenated phosphatidylethanolamines species (C18:0/C20:4 and C18:0/C22:4)^41^ and ferroptotic cell death (Fig. 3, J and K; Fig. S8G). Thus, in contrast to depleting the cysteine–GSH–GPX4 axis, ADP-ribose exposure remodels purine and pyrimidine metabolism through XO/DHODH to excessively produce superoxide from hypoxanthine and dihydroorotate, triggering initiation and propagation of lipid peroxidation and ferroptosis (Fig. S8H).

### ADP-ribose exposure induces neuronal ferroptosis *invivo*

We next evaluated whether ADP-ribose induces *in vivo* XO/DHODH-dependent neuronal ferroptosis. In *D. drosophila*, ADP-ribose feeding induced wing defects (Fig. S10A) and severely impaired climbing ability linked to neurodegeneration (Fig. S10B). ADP-ribose also significantly reduced fly survival (Fig. S10C). Consistent with our findings on ENT2 and metabolite accumulation, pretreatment with allopurinol and leflunomide before ADP-ribose extended lifespans (Fig. S10C). We overexpressed NUDT9 in TauR406W mutant flies using GAL4/UAS. NUDT9 overexpression significantly reduced brain ADP-ribose levels and prolonged lifespans (Fig. S10, D and E). Depletion of PARP1 or PARG in *D. drosophila* causes developmental arrest before pupation, therefore, we treated flies with the PAPR1 degrader iRucaparib-AP6. iRucaparib-AP6 treatment effectively blocked ADP-ribose production and extended lifespans of TauR406W mutants (Fig. S10, F and G).

Utilizing rat organotypic hippocampal slice cultures (OHSCs), which preserve the integrity of neuronal connections and supporting cells (Fig. 4A) ^42^, we found ADP-ribose treatment induced extensive neuronal cell death by diphenyl-1-pyrenylphosphine and propidium iodide staining ^43^ in the dentate gyrus (DG), CA1 and CA3 regions (Fig. 4, B to D, and Fig. S11, A to E). Attenuating XO/DHODH with AAV-knockdown or co-treating with allopurinol/leflunomide significantly reduced ADP-ribose toxicity, as did DFO and Lip-1 controls (Fig. 4, B to D, and Fig. S11, A to E).

**Fig. 4.**
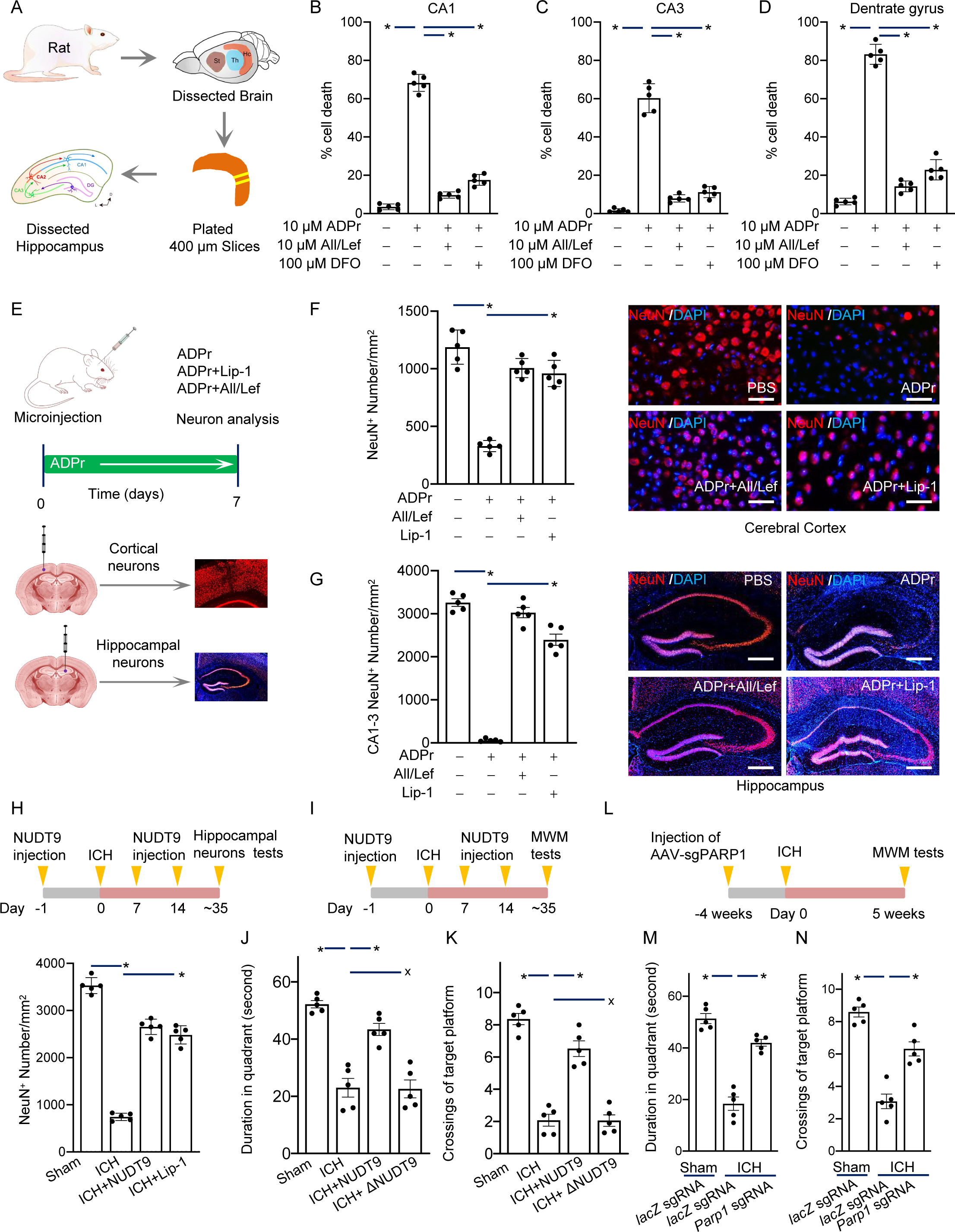
**ADP-ribose exposure induces neuronal ferroptosis *in vivo* and is responsible for ICH-induced cognitive decline in mice** A. Cartoon outline of rat organotypic hippocampal slice culture procedure. B. Organotypic hippocampal slices were pre-treated with 10 μM allopurinol/leflunomide or 100 μM DFO for 3 hours, then exposed to 10 μM ADP-ribose for 7 days. Slices were stained with the positive staining diphenyl-1-pyrenylphosphine (DPPP) and the number of positively stained cells in the CA1 region was quantified. C. Positively stained cells were quantified in the CA3 region of slices treated as in (B). D. Positively stained cells were quantified in the dentrate gyrus region of slices treated as in (B). E. Schematic diagram showing the microinjection of 1μL ADP-ribose (1mM), allopurinol/leflunomide (200μM) and Lip-1 (40μM) into the cerebral cortex and hippocampus of mice. Lip-1, liproxstatin-1, a ferroptosis inhibitor. F. Cortical neuronal damage from ADP-ribose was characterized by counting NeuN-positive cells in the cortex (n=5 mice/group). Representative microscopic fluorescence imaging of cortical neurons from 1 of 5 mice per condition. Scale bar, 40μm. G. Histogram of neuronal counts in CA1-3 regions of hippocampus from mice treated as in (E) (n=5 mice/group). Representative microscopic fluorescence imaging of hippocampal neurons from 1 of 5 mice per condition. Scale bar, 350μm. H. Hippocampal CA1 neuronal counts at 5 weeks post-ICH by measuring NeuN-positive cells. Mice received ipsilateral (relative to the collagenase-infused side) hippocampal microinjections of 1μL NUDT9 (1 µg/μL) or Lip-1 (40 μM) 1 day before ICH and on days 7/14 post-ICH. I. Schema of Morris water maze paradigm used to assess spatial learning and memory performance at 5 weeks post-ICH. Mice received ipsilateral hippocampal microinjections of 1μg NUDT9 or heat-denatured NUDT9 one day before ICH and on days 7/14 post-ICH. J. Statistical results measuring time spent in the target quadrant during the probe trial. K. Counting the times of crossing the target platform area during the probe trial. L. Schema of MWM paradigm for PARP1 knockdown ICH mice. AAV-SpCas9 (Mecp2 promoter) and AAV-SpGuide (Parp1- or lacZ(control)-targeting sgRNAs) were injected into the ipsilateral hippocampus 4 weeks before ICH (induced 1 month after AAV injection) to knock down PARP1. M. Statistical results measuring time spent in the target quadrant during the probe test. N. Counting the times of crossing the target platform area during the probe test. Data shown are means ± SD and are analyzed using two-way ANOVA (brain region × drug treatment) followed by Bonferroni posthoctests in B-D; Data shown are means ± SEM and are analyzed using Kruskal–Wallis one-way ANOVA followed by Dunn’s multiple comparison tests in F-N. n=5 mice; *, P< 0.01. See also Figures S10 to S15. Statistics source data can be found in Supplementary Table 2.

We investigated the effects of direct ADP-ribose microinjection into the mouse hippocampus and cortex (Fig. 4E). Two days post-injection, ADP-ribose severely depleted neuronal numbers in the cortex (Fig. 4F; for local magnification, see Fig. S12, A to D) and hippocampus regions (Fig. 4G; for local magnification, see Fig. S13, A to D). Specifically, most CA1-CA3 neurons died while DG neurons farther from the injection site were preserved (Fig. S13B). Critically, concurrent injection of allopurinol and leflunomide significantly attenuated ADP-ribose-mediated neurotoxicity in both the cortex (Fig. 4F; for local magnification, see Fig. S12C) and hippocampus (Fig. 4G; for local magnification, see Fig. S13C). Thus, direct ADP-ribose exposure *in vivo* induces neurological impairment by activating XO/DHODH-dependent ferroptotic signaling pathways.

### ADP-ribose is responsible for ICH-induced cognitive decline in mice

Intracerebral hemorrhage (ICH) induces oxidative stress and cognitive impairment in mice ^21,44^. We utilized a collagenase-induced ICH model to assess whether ADP-ribose is responsible for ICH-induced cognitive decline. ICH significantly increased ADP-ribose levels in hippocampal neurons 24 hours post-ICH compared to sham controls (Fig. S14, A to C) and reduced CA1 neuron numbers 5 weeks later (Fig. 4H, and S14D), indicating that ICH elicits ADP-ribose production and hippocampal neuronal death. We next assessed spatial memory via Morris water maze (MWM) five weeks post-ICH (Fig. 4I). While escape latency decreased across training days for all groups, ICH mice exhibited significantly poorer performance than sham controls each day (Fig. S14, E and F), with no difference in swim velocity (Fig. S14G). In a probe trial, sham mice spent more time in the target quadrant and crossed the platform location more often than ICH mice (Fig. 4, J and K). Thus, ICH impairs spatial memory required for the MWM task.

To determine whether inhibiting ADP-ribose production alleviates ICH-induced cognitive deficits, we knocked down PARP1 in neurons using AAV vectors. A 1:1 mixture of AAV-SpCas9 (containing neuron-specific *Mecp2* promoter) and AAV-SpGuide (*lacZ*-targeting sgRNAs as a negative control) was stereotactically injected into the hippocampus of adult male mice (Fig. S15, A and B). Co-staining 3 weeks after viral injection showed ∼85% co-transduction efficiency of the two vectors (GFP-KASH^+^/HA-Cas9^+^) in hippocampal CA1 neurons (Fig. S15C). PARP1 protein levels were decreased by 75% in the hippocampus of mice injected with *Parp1*-targeting vs *lacZ*-targeting sgRNAs (Fig. S15, D and E). We observed that ADP-ribose levels were significantly decreased in PARP1 knockdown hippocampal neurons 24 hours post-ICH (5.61 pg/cell in controls vs 1.58 pg/cell in PARP1-KD, *P*<0.0001) (Fig. S15F). In MWM training trials, PARP1 knockdown ICH mice had significantly reduced escape latencies compared to controls (Fig. S15, G to I). During the probe trial, PARP1 knockdown ICH mice spent more time in the target quadrant and crossed the platform location more often than controls (Fig. 4, L to N). Thus, inhibitin<u>g</u> ADP-ribose production improved ICH-induced spatial memory deficits.

Further, to determine whether promoting ADP-ribose degradation alleviates ICH-induced cognitive deficits, we administered intrahippocampal microinjections of recombinant human NUDT9 proteins or heat-inactivated NUDT9 before ICH and on days 7 and 14 post-ICH (Fig. 4H). NUDT9 treatment significantly increased the number of surviving CA1 hippocampal neurons 5 weeks after ICH compared to controls, consistent with the neuroprotective effects observed with the ferroptosis inhibitor Lip-1 (Fig. 4H, and Fig. S14D). In MWM training trials, ICH mice receiving NUDT9 proteins showed significantly reduced escape latencies versus those receiving heat-denatured NUDT9 (Fig. S14, E and F). During the probe trial, ICH mice receiving NUDT9 proteins spent more time in the target quadrant and crossed the platform location more frequently than controls (Fig. 4, J and K). Collectively, these results demonstrate that promoting ADP-ribose degradation through NUDT9 administration alleviated ICH-induced neuronal loss and cognitive impairment, indicating blocking ADP-ribose signaling mitigates cognitive decline from ICH.

## DISCUSSION

In this study, we demonstrate that ADP-ribose, a product of NAD^+^ consumption, induces neuronal ferroptosis and is elevated in neurological disease models such as AD and stroke. Our findings also revealed a complex interplay involving XO and DHODH in driving lipid peroxidation and subsequent neuronal ferroptosis. Interventions blocking ADP-ribose signaling alleviated cognitive decline in ICH models, highlighting the potential therapeutic significance of targeting this pathway. Therefore, a therapeutic strategy involves supplementing NAD^+^ availability while preventing ADP-ribose neuronal toxicity. To inhibit ADP-ribose accumulation, we propose three approaches: 1) inhibiting NAD^+^-consuming enzymes like PARP1 to block ADP-ribose production; 2) depleting ADP-ribose with an ADPR hydrolase like NUDT9; 3) inhibiting ADP-ribose-driven ROS generation with XO/DHODH inhibitors like allopurinol and leflunomide.

While ADP-ribose-induced neuronal cell death relies on PARP1 activation, the mechanism diverges from PAR-induced parthanatos^8^. Extracellular ADP-ribose directly binds to ENT2, remodeling intracellular purine and pyrimidine metabolism. In neurons that predominantly utilize salvage pathways for nucleosides synthesis rather than *de novo* pathways, ENT2 functions as the high-affinity transport to fulfill nucleoside demands ^45^. As ENT2 supports energy homeostasis^46^, deletion impairs mitochondrial bioenergetics and deteriorates neurological dysfunction in mice^47^. We hypothesize that in cell types resistant to ADP-ribose-induced ferroptosis, such as astrocytes, multiple nucleoside transport systems are engaged rather than sole reliance on ENT2.

DHODH contributes to mitochondrial ROS production directly and indirectly by depositing electrons onto CoQ to form CoQH_2_ that is then re-oxidized by CIII/IV-mediated respiration^48^. Importantly, DHODH inhibitors like leflunomide or brequinar bind to the CoQ site of DHODH, inhibiting both direct and indirect modes of mitochondrial ROS production^48,49^. However, the physiological relevance of DHODH-induced ROS was previously unclear. Our study found that DHODH-induced mitochondrial ROS promotes ER phosphatidylethanolamines peroxidation and neuronal ferroptosis.

While DHODH inhibition was found to reduce ADP-ribose-induced mitochondrial ROS production and lipid peroxidation in neurons, a recent study showed that DHODH protects cancer cells from ferroptosis^50^. This discrepancy may stem from the different metabolic features between cancer and neuronal cells: 1) cancer cells frequently overexpress DHODH to support rapid proliferation via *de novo* pyrimidine synthesis, whereas neurons predominantly rely on the salvage pathway for nucleic acid synthesis and DHODH is dispensable under normal physiological conditions^51,52^; 2) cancer cells maintain high redox stress signaling threshold compared to neurons, which exhibit low tolerance for oxidative stress and have high phospholipid content in membranes, enhancing vulnerability to lipid peroxidation and ferroptosis ^53^; 3) neurons also appear to be intolerant of glutamine^54^, which is essential for *de novo* pyrimidine synthesis in cancer cells; 4) there may be off-target effects of DHODH inhibitors not directly linked to DHODH itself^55^. Our study reveals that distinct metabolic features determine cellular responses to ferroptotic insults.

### Limitations of the Study

This study provides novel insights into the mechanisms of ADP-ribose signaling and neuronal ferroptosis. However, there are some limitations that future work could address. First, our findings that ADP-ribose induces neuronal cell death relies heavily on *in vitro* studies using cultured cells and acute toxin exposures. Validating conclusions regarding ADP-ribose signaling and their perturbation in rigorously characterized *in vivo* disease paradigms will strengthen the pathophysiological relevance of our findings. Additionally, our findings suggest ADP-ribose production transcends PARPs, as oxidatively stressed neurons release ADP-ribose independently PARP1. Precisely characterizing how ADP-ribose production relates to specific NAD^+^-consuming enzymes, such as Sirtuins and SARM1, under different disease contexts remains an important area for future study. Lastly, while our study focused on extracellular ADP-ribose functions in neurons, TPP screening identified 40 intracellular proteins whose stability was altered by ADP-ribose binding. Elucidating the spatio-temporal regulation and functional consequences of intracellular ADP-ribose engagement with putative targets in neurons and across diverse cell types, could provide novel mechanistic insights. Overall, continued interrogation of ADP-ribose signaling in health and disease holds promise for advancing our understanding and ability to intervene against neurological disorders.

## ACKNOWLEDGMENTS

We thank Hamid Baniasadi (UT Southwestern Medical Center) for untargeted metabolomics and target metabolomics analysis, Xiaoyu Wang (UT Southwestern Medical Center) for ADP-ribose concentration analysis, Duojia Pan (UT Southwestern Medical Center) for feeding of Drosophila, and Wei Peng (UT Southwestern Medical Center) for protein-molecule structure mock docking. We thank Marcus Conrad (Helmholtz Zentrum München), Boyi Gan (University of Texas MD Anderson Cancer Center), Xuejun Jiang (Sloan Kettering Institute), Yonghao Yu (UT Southwestern Medical Center), Baiyun Wang (University of Washington) and Jiang Dai (Jingwan Biomedical Technology Co., LTD) for helpful discussions and suggestions.

## Author contributions

L.Y. conceived the study, designed, performed, and interpreted the experiments and wrote the manuscript. Most of the experiments were performed by L.Y., H.N., Y.H. and H.C.; P.C., G.L. and B.C. developed and performed bioinformatics and statistical analysis, and revised the manuscript; H.L., P.C., and F.Y. generated the fruit fly model and mouse model, and revised the manuscript; AD patient sample collection and analysis was carried out by Q.Z., H.N., B.C. and L.Y..

## Supplementary Tables

Supplementary Tables 1–8.

## METHODS

### Ethics statement

All animal experiments were conducted in accordance with the ‘Guide for the Care and Use of Laboratory Animals’ and ‘Principles for the Utilization and Care of Vertebrate Animals’, and approved by the Institutional Human Research Ethics and Animal Care and Use Committee at Wenzhou Medical University, Peking University and Peking University Shenzhen Hospital. 2×FAD and 5×FAD mice were from Yong Zhang (Peking university). We obtained human aged 55 to 70 years old lumbar CSF from the sample bank of the department of neurology, at Peking University Shenzhen Hospital and First Affiliated Hospital of Wenzhou Medical University. Diagnosis of Alzheimer’s disease was established according to the NINCDS-ADRDA criteria. In all patients and controls, lumbar puncture was performed, and CSF Chromogranin A measured. Control subjects underwent CSF examination due to lower back pain or a tension-type headache. The study protocol, including CSF examination, was approved by the Ethics Committee of Peking University Shenzhen Hospital (# 2017S3-036). Informed consent was obtained from all subjects or their relatives, and no compensation was provided for the sample collection.

### Cell culture, compounds and antibodies

SH-SY5Y and HEK293T cells were purchased from ATCC, and HT22 cells were purchased from Sigma. All cell lines were authenticated by STR profiling and tested for mycoplasma contamination by GENEWIZ. Cells were cultured in DMEM supplemented with 10% FBS. Cells were transfected with various plasmids using Lipofectamine 3000 (Invitrogen) reagent according to the manufacturer’s protocol. To provide a more intuitive and comprehensive overview of the key resources utilized in this study, including compounds, plasmids, and antibodies, we have included a detailed key resources table in the attached document named Supplementary_Table_6_KEY RESOURCES TABLE.

### Isolation and culture of primary cortical neurons

Primary cortical neurons were isolated from embryonic day 16.5 (E16.5) mice. The cortex without meninges was carefully extracted from brain in the ice-cold HBSS and dissociated into small pieces. We collected dissociated cortex through centrifugation for 3.5min at 500g. Then the shredded cortex tissues were suspended and incubated in Enzyme solution (HBSS + 10U/mL papain + 1mM CaCl_2_ + 0.5 mM EDTA+ 1% DNase) for 10-15min with occasional mixing at 37℃. Then the digestion was ceased by 10% FBS and the cells were collected by centrifugation.

The single cells suspension was resuspended by 1mL DMEM and harvest by pipetting the digested cortex tissue for 10 times and passing a 40 μm filter. Cells was harvest again, resuspended by neurobasal medium (Gibco, USA) with 1% FBS and plated into PDL-coated 24-wells plates (5 × 10^5^/well). Three hours later, the medium was changed into neurobasal medium with 2mM GlutaMAX (with or without 5μM ARAC, according to gestational day of mice; Gibco, USA) but no FBS.

Neurons were maintained at 37°C with 5% CO_2_ for 24 hours after plating prior to experiments using immature primary cortical neurons. The neurons were validated through visual morphology and positive expression of the neuronal marker NeuN. These immature cultures are sensitive to 5mM glutamate/L-homocysteic acid (HCA)-induced ferroptosis. This cell death is independent of ionotropic glutmate receptors, which do not fully develop until much later *in vitro* (> 10 days). HCA-induced death in immature neuronal cultures is insensitive to MK801 (an ionic glutamate receptor antagonist; Sigma Aldrich) but is rescued by supplemental cyst(e)ine.

For mature primary cortical neuronal cultures, every three days 50% media was changed and cells were allowed to mature for 14 days before experimentation. Neurons were identified through visual morphology. Mature neurons were sensitive to glutamate-induced excitotoxicity as cell death was completely inhibited by MK801 treatment. Excitotoxicity was induced by treating mature neurons with 100μM glutamate (Sigma Aldrich) in HBSS buffer for 1 hour at 37°C, followed by restoring the media at 37°C for 24 hours. The glutamate receptor antagonist MK801 (10μM; Sigma Aldrich) was used as a positive control and added immediately after glutamate treatment. Cell viability was assessed using the MTT assay (Promega).

### Drosophila culture and strains

The transgenic flies were constructed to bear the different human tau constructs (tau, tauR406W) under the control of a yeast upstream activating sequence (UAS). The human tau constructs were cloned into the pUAST vector for the Gal4-inducible expression of transgenic pseuhosphorylated tau in flies. Transgenic flies were generated by BestGene. Flies with the following genotypes were utilized: 2N4R tau (UAS-tau/CyO) and 2N4R R406W (UAS-tauR406W/CyO). The expression of tau protein was spatial controlled crossing transgenic flies with two driver lines with specific expression to the nervous system (Elav-Gal4/CyO, Bloomington Stock Centre, Indiana, USA, stock numbers 8765 and 25750). Only the progeny without curly phenotype was analyzed (Elav-Gal4/UAS-tau; Elav-Gal4/UAS-tauR406W). All Drosophila stock and crosses were reared on standard Drosophila media of sugar-cornmeal-yeast-agar and maintained at 23℃. To induce the expression of the human tau, flies were raised at 29℃.

### CSF Sample Preparation

After successful administration of spinal anesthesia, but prior to the administration of local anesthetic, CSF (2 mL) was collected in a polypropylene tube and placed on ice. Samples were immediately centrifuged at 3,000 × g for 10 min at 4°C to remove cells (Muller et al., 2014), and the supernatant was aliquoted and stored at –80°C until analysis. Metabolites and lipids were extracted from CSF samples using liquid–liquid extraction. Briefly, 500 μL of CSF were concentrated to 50 μL and mixed with a fourfold volume of ice-cold chloroform/methanol (2:1, v/v). After vertexing for 15 min at 4°C, the mixture was centrifuged at 13,000 × g for 15 min. The upper aqueous phase (hydrophilic metabolites) and lower organic phase (hydrophobic metabolites) were separately collected and evaporated at room temperature under vacuum. All evaporated samples were stored at −80°C until high-performance liquid chromatography–mass spectrometry (HPLC–MS) analysis.

### *In vitro* model of ferroptosis in neurons

Cell death was induced in primary cortical neurons and hippocampal HT22 cells by treating with 100 μM hemin (Sigma Aldrich). The stock solution for hemin was dissolved in 0.1M NaOH. Primary cortical neurons and HT22 hippocampal neuroblasts were exposed to 20 μM and and 2 μM erastin (dissolved in DMSO; Selleck Chemicals), respectively, to induce oxidative stress-induced cell death by blocking xc^-^ transporters. Primary cortical neurons and HT22 hippocampal neuroblast cells were exposed to 5mM L-homocysteic acid (HCA; Sigma Aldrich) dissolved in MEM no phenol (Thermo Fisher Scientific), a structural analog of glutamate, to induce oxidative stress – induced cell death by blocking the xc^-^ transporter. Sodium selenite (Sigma Aldrich), dissolved in ddH_2_O was applied to the media at varying concentrations. The time of addition of hemin, erastin or HCA varied depending on experimental design; 24 hours after HCA treatment, the cells were rinsed with warm PBS and cell viability was assessed by MTT assay (Promega) or calcein-AM/ethidium homodimer-1 staining (Live/Dead assay, Thermo Fisher Scientific). Similarly, other known ferroptosis inducers, RSL3 (Cayman Chemical) or FIN56 (Sigma Aldrich) dissolved in DMSO, were used to induce cell death in primary cortical neurons. A minimum of 3 independent cell culture experiments were performed with studies involving ferroptosis of cells *in vitro*.

### ADP-ribose concentration measurement

**Analytical Processing**: 1×10^7^ cells were harvested and washed 3 times with PBS. Add 1 ml of 80% methanol (pre-cooled at -20°C), shake well and centrifuge at high speed to remove protein precipitates. Aliquot 500 μL 80% MeOH for blank and 490 μL 81.63% MeOH for standards. Spike standards with appropriate amount of H_2_O stock. Vortex 15 second. Transfer 200 μL of blank and standards to the second Eppendorf tube. Aliquot 200 μL samples to the Eppendorf tubes. Add 1 μL (10 ng/μL) of IS solution (IS final conc. = 50 ng/mL) to each sample including blank, standards and samples, Vortex. Take 180 μL supernatant and transfer to HPLC vial with insert. The supernatant was then analyzed by LC/MS/MS. **QTrap 6500 Parameters**: Method: ADPr + pHilic UMP-13 IS 010920; Ion Source/Gas Parameters: CUR = 45, CAD = High, IS = 2000, TEM = 500, GS1 = 70, GS2 = 70; Buffer A: dH2O + 20 mM NH4AC, Buffer B: 95%ACN +5% dH2O + 2 mM NH_4_AC+ 0.1% HAC; flow rate 0.25 ml/min; column: Zic pHillic PEEK coated 150 x2.1 mm 5 μM column; 0-1.5 min 80% B, 1.5-5.0 min gradient to 30% B, 5.0-8.0 min 30% B, 8.0 - 9.0 min gradient to 80% B, 9.0 - 12.0 min 80% B; Compound transition 559.895 to 347.800; IS UMP-13 transition 336.092 to 102.000. **Explaining**: intracellular ADP- ribose concentration (pg/cell) = total ADP-ribose mass in living cells /total living cell numbers; extracellular ADP-ribose concentration (pg/cell) = total ADP-ribose mass in cell culture medium/total cell numbers; ADP-ribose concentration in cerebrospinal fluid (CSF) (μg/ml) = total ADP-ribose mass in CSF/ CSF volume.

### Determination of intracellular metabolite content

Aspirate the medium and wash cells 2-3 times with PBS. Then add PBS, scrape the adherent cells with a spatula, and count the cells. Take about 1 × 10^7^ cells, centrifuge to remove the supernatant. Add 1 ml of methanol/water (4:1) mixture (pre-cooled at -20°C), shake well and centrifuge at high speed to remove protein precipitates. Centrifuge again and transfer the supernatant to a new EP tube. The intracellular metabolites were measured by LC-MS QTrap 6500.

### Organotypic Hippocampal Slice Cultures

OHSCs were cultured as previously described^56^ with approval from Wenzhou Medical University’s IACUC. Briefly, Sprague Dawley rat pups (P6-P7) were rapidly decapitated, and the hippocampus placed in ice-cold Hanks’ Balanced Salt Solution (Gibco). 400 μm thick sections were cut using a McIlwain tissue chopper and immediately plated on 0.4-mm pore size Millicell cell culture membrane inserts (Millipore) in 800μL MEM culture Medium, containing: heat-inactivated horse serum 20%, HEPES 30mM, glutamine 2mM, D-glucose 13mM, NaHCO_3_ 5mM, CaCl_2_ 1mM, MgSO_4_ 2mM, insulin 4 mg/L at pH 7.2 and osmolarity 320. Cultured slices were maintained at 37°C and 5% CO_2_. Medium was changed every 1-2 days.

### Excitotoxic Injury on OHSCs

After 10-14 DIV, OHSCs were exposed to serum free media (SFM) consisting of 10 μM ADP-ribose for 24 hr. Only healthy OHSCs defined as those with less than 5% cell death in all regions (DG, CA3, CA1) of the hippocampus pre-injury were used for experiments. If chemical compounds (*e.g.*, allopurinol/leflunomide and DFO) were added, they were added 3 hr before the ADP-ribose.

### Analysis of Cell Death in Rat Brain Slices

Quantification of cell death has been described previously^9^. Brightfield images of the hippocampal cultures were taken before injury for identification of regions of interest (ROI) including the dentate gyrus (DG), CA3 and CA1. Diphenyl-1-pyrenylphosphine (DPPP, SANTA CRUZ BIOTECHNOLOGY) was used as a fluorescent signal for lipid peroxidation (ferroptosis mark), and images were taken before the induction of injury and 24 hr following injury. For DPPP/PI imaging, the cultures were transferred to SFM supplemented with 2 μM DPPP/PI. After a 30min incubation, brightfield and DPPP/PI images were acquired. All images were captured on a Leica DMi8 fluorescent microscope. Metamorphic image analysis software was used to determine the ROI in the brightfield image, and this ROI was transferred to the DPPP/PI image taken before and 24 hr after injury. Percentage cell death was expressed as the number of pixels in the ROI above a threshold in the DPPP fluorescent image divided by the total number of pixels in the ROI.

### Analysis of 4-HNE Production

Lipid Peroxidation (4-HNE) Assay Kit (ab238538) is used for the quantitation of 4-HNE adduct in protein samples. The standard curve is generated with 4-HNE-BSA. 50 μL standard or sample were added to wells of 4-HNE Conjugate coated plate and incubated for 10 mins. Add 50 μL of the diluted anti-4-HNE antibody and incubate for 1 h. Washing steps with 250 μL 1×Wash Buffer. Add 100 μL diluted Secondary Antibody-HRP Conjugate per well and incubate for 1 h. Wash as before with 1×Wash buffer. Add 100 μL of warm Substrate Solution and incubate for 20 mins. Stop the enzyme reaction by adding 100 μL of Stop Solution to each well. Read absorbance immediately on a microplate reader using 450 nm.

### Analysis of Reactive Oxygen Species Production

The day before the experiment, 5 × 10^5^ cells/well were seeded in 6-well dishes (Corning). The day of the experiment, cells were treated with test compounds for the indicated times; harvested by trypsinization; resuspended in 500 μL Hanks Balanced Salt Solution (HBSS, Gibco) containing H_2_DCFDA (from abcam, ab113851); and incubated for 30 min at 37°C in a tissue culture incubator. Cells were then resuspended in 500 μL buffer, and analyzed using a microplate reader by fluorescence spectroscopy with excitation / emission at 485 nm/535 nm.

### Lipid Peroxidation Measurement

Cells were seeded on 12-well plates and incubated overnight. The next day, cells were treated with compounds for the indicated times, harvested by trypsinization and resuspended in 50 μL lysis buffer (ab118970). Cell lysates were incubated with 10 μL MDA Color Reagent (ab233471) for 30 min at 25°C in a water bath. Add 40 μL of Reaction Solution and incubate at room temperature for 60 minutes. Monitor absorbance increase at 695 nm.

### Superoxide Anion Assay

The method is based on the oxidation of luminol by superoxide anions resulting in the formation of chemiluminescence light. This method utilizes a specific, non-toxic enhancer that amplifies the chemiluminescent signal. Resuspend the cells in the Assay Medium (Sigma, A6105) to a concentration of 5.0×10^5^ to 1.0×10^6^ cells/100 μL. Start the reaction by adding to each well 100 μL of Assay Buffer (Sigma, A5980) containing with 5% Luminol Solution (Sigma, L5034). After 30min, measure the luminescence intensity by end point method.

### Identification of PE oxygenated products in cell lysates

SH-SY5Y cells pre-treated with 10μM allopurinol and leflunomide for 1 hour were treated with 100μM ADP-ribose for 12 hours. Subsequently, the cells were resuspended in 25 mM HEPES buffer (pH 7.4) containing 200 μM DTPA, sonicated, and prepared for experimentation. Lipids were extracted using the Folch procedure. The PE fraction, containing both non-oxygenated and oxygenated PE was obtained from those cells by normal phase LC/MS. PE and PEox (PEOOH) were separated on a reverse phase column (Luna 3 μm C8 (2) 100A, 150 × 4.6 mm, Phenomenex). The column was maintained at room temperature and eluted at a flow rate of 1.5 ml/min using an isocratic solvent system consisting of acetonitrile/H2O/trimethylamine/ acetic acid (45/5/0.5/0.5, v/v/v/v). All solvents were of LC/MS grade. The eluant was monitored by UV absorbance at 205 and 235nm on a Shimadzu HPLC system. Fractions containing PEox were collected and identification was confirmed by mass spectrometry.

Oxygenated PEs were identified by MS/MS analysis utilizing an Orbitrap Fusion Lumos mass spectrometer (ThermoFisher Scientific). The instrument was operated with an electrospray ionization probe in negative polarity mode, with the following ion source conditions: spray voltage = 3 kV, sheath gas = 55 (arbitrary unit), auxiliary gas = 10 (arb. unit), sweep gas = 0.5 (arb. unit), transfer tube temperature = 300 °C, vaporizer temperature = 200 °C, RF-Lens level = 20 %. Data acquisition was performed in data-dependant-MS^2^ targeted-MS^3^ mode, with a cycle time setting of 3 s. For the MS scan event, the parameters were set as follow: ion detection using the Orbitrap, mass resolution of 120,000, scan range of m/z 400 – 1800, and AGC target of 1e5. The most intense ion was selected for the data-dependent MS^2^ scan, with a dynamic exclusion of 9 s. An exclusion mass list for MS^2^, consisting of m/z values of 130 background ions, was created from solvent blank injection data. For the MS^2^ scan event(s), the parameters were as follows: quadrupole isolation of 1 Da, first mass of m/z 87, activation type of HCD, collision energy of 28 % with a step of 8 %, ion detection using the Orbitrap, mass resolution of 15,000, max injection time of 250 ms, and AGC target of 2e4. Product ions from a targeted mass list were selected for MS^3^ scanning. The target mass list for MS^3^: 319.2279, 317.2123, 335.2228, 333.2072, 317.2123, 351.2159, 333.2054, 349.2003, 331.1898, 367.2126, 349.2021, 331.1916.

For the MS^3^ scan event(s), the parameters were set as follows: top N = 4, isolation window = 2 Da, activation type = HCD, collision energy = 40 %, ion detection = Ion Trap, ion trap scan rate = Rapid, max injection time = 500 ms, and AGC target = 3e4.

Normal phase column separation and MS/MS analysis of phospholipids Phospholipids were separated on a normal phase column (Luna 3 μm Silica (2) 100Å, 150 × 2.0 mm, Phenomenex) at a flow rate of 0.2 mL/min on a Dionex Ultimate 3000 HPLC system. The column was maintained at 35°C. Gradient elution was achieved using solvents A and B, both containing 10 mM ammonium acetate and 0.5% triethylamine. Solvent A comprised propanol:hexane:water (285:215:5, v/v/v) and solvent B sonsisted of propanol:hexane:water (285:215:40, v/v/v), all of LC/MS grade. The elution profile began with an isocratic hold at 25% B for 0.5 min, follwed by a linear gradient of 25–40% solvent B from 0.5 to 6.5 min, 40–55% solvent B from 6.5–25 min, 55–70% solvent B from 25–38 min, 70%–100% solvent B from 38– 48 min. The column was then held isocratically at 100% solvent B for 7 min before returning to initial conditions (25% B) over 15 min. The column was equilibrated at 25% B for an additional 5 min.

MS and MS/MS analysis of PLs was performed on a Q-Exactive hybrid-quadrupole-orbitrap mass spectrometer (ThermoFisher Scientific) in negative ion mode. Full MS scans were acquired at a resolution of 140,000, while MS^2^ scan were acquired at a resolution of 17,500 in a data-dependent manner. The MS scan range was set to 400–1800 m/z with a maximum injection time of 128 ms using 1 microscan. For MS^2^ analysis (high energy collisional dissociation (HCD)), a maximum injection time of 500 ms was used, with a collision energy set to 24. An inclusion list for phospholipids, including PE, PC, and CL, as well as their oxidized and deuterated products, was employed. An isolation window of 1.0 Da was set for both MS and MS^2^ scans. The capillary spray voltage was set to 3.5 kV, and the capillary temperature was maintained at 320 °C. The S-lens Rf level was adjusted to 60.

### Glutathione Assay

Collect cells by centrifugation (i.e., 1,000-2,000 x g for 10 minutes at 4°C). The cell pellet can be homogenized or sonicated in 1-2 mL of cold buffer (i.e., 50 mM MES or phosphate, pH 6-7, containing 1 mM EDTA). Centrifuge at 10,000 x g for 15 minutes at 4°C. Remove the supernatant and store on ice. Add an equal volume of the 1g/mL MPA Reagent to the sample and mix by vertexing. Allow the mixture to stand at room temperature for five minutes and centrifuge at >2,000 g for at least two minutes (a microfuge will be sufficient for the centrifugation). Carefully collect the supernatant without disturbing the precipitate. Add 50 μL of 4M TEAM Reagent per ml of the supernatant and vortex immediately. The TEAM Reagent will increase the pH of the sample. Read the plate at 405-414 nm after 25 minutes.

### ^13^C-tracers Uptake Assay

SH-SY5Y cells were seeded in 6-well dishes (Corning) at a density of 5 × 10^5^ cells per well. The next day, cells were treated with 100 μM ADP-ribose for 3 hours; cells were washed twice in prewarmed uptake buffer (137 mM choline chloride, 3 mM KCl, 1 mM CaCl_2_, 1 mM MgCl_2_, 5 mM D-glucose, 0.7 mM K_2_HPO_4_, and 10 mM HEPES pH 7.4) and then incubated for 10 min at 37°C in 1 mL of uptake buffer to deplete cellular amino acids. At this point, in each well, the buffer was replaced with 600 μL uptake buffer containing ADP-ribose and 50 μM of ^13^C-tracers (uridine cytidine, adenosine and cystine) and incubated for 3 min at 37°C. Cells were then washed three times with ice-cold uptake buffer and lysed in 500 μL 0.1 M NaOH. This lysate was measured by LC-MS QTrap 6500. All measurements were performed in triplicate for each condition.

### Neuronal Cell Viability Measurements

Cell viability was typically assessed in 96-well format by Cell Counting Kit-8 (CCK-8, Dojindo) and alamarBlue^TM^ Cell Viability Reagent Blue. Briefly, cells were seeded onto 96-well plates at a density of 2×10^4^ per well. Subsequently, cells exposed to 10 μl CCK-8 reagent (100 μl medium per well) for 1 hour at 37 °C, 5% CO_2_ in an incubator. The absorbance at a wavelength of 450 nm was determined using a FLUOstar Omega microplate reader (BMG Labtech). Alamar Blue (Invitrogen) fluorescence (ex/em 530/590) measured on a Victor3 plate reader (Perkin Elmer). In some experiments, Trypan blue dye exclusion counting was performed by using an automated cell counter (ViCell, Beckman-Coulter). Cell viability under test conditions is reported as a percentage relative to the negative control treatment.

### Iron Ion Analysis

The ferrous and total iron was quantified using a spectrophotometric iron assay kit (BioVision, Mountain View, CA) according to the manufacturer’s directions. To quantify iron, samples were treated with a reducing agent prior to incubation with Ferene S (Sigma-Aldrich, St. Louis, MO). The optical density at 593 nm (OD_593_) was measured after 1 h with a Synergy 2 microplate reader (Biotek, Winooski, VT). A standard curve was routinely established (0 to 10 nmol of iron).

### ATP Concentration Analysis

Cells were lysed in RIPA lysis buffer (50mM Tris-HCl pH 7.4, 150mM NaCl, 1% Nonidet-P40, 0.5% sodium deoxycholate, 1% SDS). After 20 min centrifugation at 12,000 rpm at 4 °C, the supernatant was collected. The ATP concentration of the supernatant was assessed quantitatively with the EnzyLight™ ATP Assay Kit (BioAssay Systems, Hayward, CA) according to the instructions.

### Thermal Proteome Profiling (TPP)

Lysis (M-PER buffer, Thermo Scientific, 89842 and 78501) was performed by five freeze–thaw cycles using a 25 °C heating block and liquid nitrogen. Cell debris and precipitated proteins were removed by centrifugation at 21,000g and 4 °C for 40 minutes. Supernatants were transferred to new tubes and incubated with compounds at room temperature for 10 minutes then placed on ice for 10 minutes. Equal volumes of soluble protein supernatants were transferred to new tubes and subjected to gradient temperature denaturation for 3 minutes. Precipitated proteins were removed by centrifugation at 21,000g and 4 °C for 40 minutes. First, the supernatants samples were supplemented with reagents to contain a final concentration of 50 mM TEAB, 0.1% SDS and 5 mM TCEP. Reduction was performed at 65 °C for 30 minutes. Samples were then cooled down to room temperature and alkylated with 15 mM of chloroacetamide for 30 minutes. Proteins were digested overnight with 1:40 Lys-C (Wako Chemicals)-to-protein ratio. Consecutively, trypsin (Thermo Fisher Scientific) was added at a 1:70 enzyme-to-protein ratio for an 8-hour incubation at 37 °C. Finally, the same amount of trypsin was added one more time for an overnight incubation. Resulting peptides were labeled by 10-plex TMTpro tags (TMTpro, Thermo Fisher Scientific) using the same amount of respective label for each sample. Eight melting points of two randomly selected cell lines were combined in each TMT 10-plex set. The protein amounts were adjusted to contain the same total protein amount for all cell lines throughout the TMT sets. An overview of the sets is given in Supplementary Table 1. Labeling was performed according to the manufacturer’s instructions but with 2-hour incubation before quenching the TMT labeling reaction. Labeling efficiency was determined by LC–MS/MS before mixing the TMT-labeled samples. Sample cleanup was performed using solid-phase extraction Strata-X-C SPE columns (Phenomenex). Purified peptides were dried in a vacuum centrifuge.

### RNA preparation and quantitative real-time PCR analysis

Different mouse tissues or cells were lysed in Trizol (Invitrogen). Total RNA was recovered from tissues following the manufacturer’s protocol. Two micrograms of purified RNA from each sample were reverse transcribed to single-stranded cDNA with All-In-One RT MasterMix (catalogue no. G486, abm) The newly synthesized cDNA was mixed with TransStart Top Green qPCR SuperMix (catalogue no.AQ131, Transgen Biotech) in a volume of 20 μL. For Quantitative PCR, the Real Time PCR Detection system (ABI 7500) was using to detect each gene in triplicate. Fold changes were analyzed (quantified) relative to the internal control gene GAPDH based on the 2^−ΔΔCT^ Method. Primer sequences used for qPCR are listed in Supplementary Table 7.

### Lentivirus infection

shRNA lentiviral vectors targeting human ENT1 (TRCN0000043643, TRCN0000043645, TRCN0000043647) and ENT2 (TRCN0000043658, TRCN0000043659, TRCN0000043660) were from Sigma. shRNA lentiviral vectors targeting mouse PARP1 (sc-29438-SH) and PARG (sc-152026-SH), human XO (sc-41692-SH) and DHODH (sc-77142-SH) were from SANTA CRUZ BIOTECHNOLOGY, INC.. Lentiviruses carrying overexpressing lentiviral vectors were constructed by ourselves. The viruses were used to infect cells in the presence of Polybrene. Forty-eight hours later, SH-SY5Y cells were cultured in medium containing puromycin for the selection of stable clones. The clones stably knocking down ENT1/ENT2/XO/DHODH were identified and verified by qPCR.

### Production of concentrated AAV vectors

Adeno-associated viral (AAV) vectors for SpCas9 and guide RNAs were from Feng Zhang’s Lab. Adeno-associated virus carrying SpGuide vectors targeting mouse Parp1 (5′-ACCCTGAAAGGGTAGCTTGT-3′), Ent1 (5′-CGACTACGTCTTTGCTCAAC-3′), Xo (5′-CCAGCATGCAACGTACAGGG-3′), Dhodh (5′-TTGATCCAGAGTCGGCGCAC-3′) and negative control LacZ (5′-TGCGAATACGCCCACGCGAT-3′) were constructed by ourselves.

High titer AAV-PHP.eB particles were produced using 1:2 ratio AAV-PHP.eB serotype plasmids and pAdDF6 helper plasmid in HEK293FT and purified on heparin affinity column. Briefly, HEK293FT cells were transfected with transgene plasmid, pUCmini-iCAP-PHP.eB and pAd-deltaF6 (pAdDF6) at 1:2:1 ratio using PEI “MAX”. Culture medium was collected after 48 h and filtered through a 0.45 μm PVDF filter (Millipore). Tittering of viral particles was done with qPCR.

### Adeno-associated viral infection

Mice were anesthetized by intraperitoneal (i.p.) injection of 100 mg/kg ketamine and 10 mg/kg xylazine. Preemptive analgesia was given (Buprenex, 1 mg/kg, i.p.). Craniotomy was performed according to approved procedures, and 1 μL of 1:1 AAV mixture (1 × 10^13^ vector genomes (Vg)/ml) of sMecp2-SpCas9; 6 × 10^12^ Vg/ml of Parp1-sgRNA;) was stereotactically injected into the hippocampus of adult mice with a coordinate of anterior/posterior, -2.0 mm; medial/lateral, ±1.4 mm; and dorsal/ventral from skull, -1.7 mm. The skull was thinned using a dremel drill with occasional cooling with saline, and the remaining dura was punctured using a glass micropipette filled with the virus suspended in mineral oil. After each injection, the pipette was held in place for 5 min before retraction to prevent leakage. The incision was sutured and 500 μL of saline was injected subcutaneously to each mouse to avoid dehydration. Proper post-operative analgesics (Meloxicam, 1–2 mg/kg) were administered for 3 d following surgery.

### Light Microscopy

Phase contrast images were acquired using an AMG EVOS FL (Advanced Microscopy Group) microscope equipped with a 10 × phase-contrast objective. Three independent fields were acquired for each experimental condition. Representative samples from one field of view are shown.

### Flow cytometry

Cells fixed overnight with cold 70% ethanol were digested with RNase at 37℃ for 30 min and stained with P.I., followed by flow cytometry analysis with a BD FACSVerse^TM^ instrument.

### Collagenase-induced mouse model of ICH

All surgeries were conducted under aseptic conditions by a skilled animal surgeon. ICH was induced by intracerebral injection of collagenase in male C57BL/6 mice (10 to 12 weeks of age). Female mice were not used to minimize the influence of sex steroids on recovery. Mice were anesthetized with isoflurane (2 to 5%) and placed on a stereotaxic frame. During the procedure, the animal’s body temperature was maintained at 37 ℃ with a homeothermic blanket. Using a nanomite syringe pump (Harvard Apparatus) and a Hamilton syringe, 1 μL of collagenase (0.05 U; Sigma) was infused into the right basal ganglia at a flow rate of 0.1 μL/min. The injection coordinates were 2.3 mm lateral to midline and 0.5 mm anterior to bregma, and the needle was inserted to a depth of 3.7 mm beneath the skull. In control animals, 1 μL of saline was infused. Following infusion, the needle was removed after a 5 min pause to minimize overflow. The burr hole was closed with bone wax, and the incision was sutured after surgery. In total, 500 μL of saline was injected subcutaneously to each mouse to avoid dehydration.

### Animals and ADP-ribose injections

Wild-type C57BL/6 mice were purchased from the Shanghai Laboratory Animal Center (Shanghai, China). Both male and female mice at 2–4 months of age were used for experiment unless otherwise stated. Mice were kept under a controlled temperature and a 12h light, 12h dark cycle with free access to water and food in the animal facility of Wenzhou Medical University. All experimental procedures and protocols were approved by the Animal Use and Care Committee of Wenzhou Medical University. Adult mice were anesthetized by a cocktail of ketamine (1.2 mg/kg body weight) − xylazine (0.8 mg/kg body weight) and placed on stereotaxic frame (KOPF, California, USA). A final volume of 1μL ADP-ribose solution with a concentration of 1mM was stereotactically injected into the hippocampus of adult mice with a coordinate of anterior/posterior, -2.0 mm; medial/lateral, ±1.4 mm; and dorsal/ventral from skull, -1.7 mm or into cortex with a coordinate of anterior/posterior, -2.0 mm; medial/lateral, ±1.4 mm; and dorsal/ventral from skull, -1 mm.

### Hematoma Volume Measurement

Mice were sacrificed 24h after ICH; the brains were removed and flash frozen in Freeze’ IT (Fisher Scientific). Coronal sections were sliced at 30 μm and placed directly on a glass slide. To quantify hematoma volume, sections were digitized at standardized coronal levels. A blinded user would measure the contours and hematoma from left and right hemisphere from each section. Hematoma volume and swelling were measured and calculated using Axiovision software (Carl Zeiss). Hematoma measurements were conducted on at least 5 independent brains per group and hematoma volume was measured throughout the brain.

### Morris water maze assessment

Cognitive function was evaluated using the Morris water maze (MWM). This test was conducted in 2 periods. During the training period, mice introduced into a circular pool could escape by finding a submerged hidden platform in the northeast quadrant. In each trial, mice were introduced to the pool through south or west sides and allowed to swim a maximum of 120 seconds to find the platform. Mice that successfully found the platform within 120 seconds were allowed to stay on the platform for 10 seconds, whereas mice that failed to locate the platform were placed on the platform and were also allowed to stay for 10 seconds. The training was conducted in 3 trials per session, with 1 sessions/d for 5 days. The swimming velocity, escape latency, and total distance swum during a trial were recorded. The water temperature in the pool was maintained at 24°C during the test. On the sixth day, the platform was removed, and a single trial was conducted by introducing the mice to the pool through the southwest quadrant. Each mouse was allowed to swim for 120 seconds, during which the time spent in the target quadrant and crossing times of the target platform were recorded. An EthoVision 3.1 tracking system (Noldus Information Technology, Wageningen) was used for recording and analyzing data.

### Immunohistochemistry

Mice were euthanized by CO_2_ overdose and perfused with intracardial injection of ice-cold PBS and 4% PFA (paraformaldehyde) in PBS. Brains were collected and postfixed with 4% PFA for 48 hr. Postfixed brains were cryoprotected with 30% sucrose solution in PBS for 72 hr and cut into 18-μm-thick sections with a cryostat. Brain sections were serially collected and stored at -20 ℃ . The primary antibody NeuN (Abcam, ab177487, 1:5000) was used for immunohistochemical analyses. Alexa Fluor 555mm or 647mm secondary antibodies from Jackson ImmunoResearch Laboratories were used at 1:5000 dilution for fluorescence. Nuclei were counterstained with DAPI. Images were captured and examined by using a Leica DMi8 fluorescent microscope. ImageJ was used for cell counting.

### Statistics and reproducibility

Experimental data were analyzed and processed using Excel (2016) and plotted with GraphPad Prism 9.4.0 (GraphPad Software). Statistical analyses were conducted using unpaired two-tailed t-tests and one-way ANOVA with Dunnett correction for multiple comparisons. For comparisons between two groups, either an unpaired two-tailed *t*-test or Mann–Whitney test was employed. For comparisons involving more than two groups, Kruskal–Wallis one-way ANOVA followed by Dunn’s multiple comparison tests or a two-way ANOVA followed by Bonferroni *post hoc* tests was utilized. Survival curves were derived from Kaplan–Meier estimates and compared using log-rank tests. All results are shown as the mean ± s.e.m. of multiple independent experiments, not technical replicates. The number of experiments, sample size and statistic tests used are detailed in the respective figure legends. Sample size was not predetermined using any statistical method. Animals were randomly assigned to experimental groups to ensure unbiased group allocation. The order of experimental conditions or stimulus presentation was not randomized. No data were excluded from the analyses. The investigators were not blinded to group allocation during experiments and outcome assessment. All statistical tests were conducted on a two-sided basis, with *P* values <0.05 considered statistically significant.

## Reporting summary

Further information on research design is available in the Nature Portfolio Reporting Summary linked to this article.

## Data availability

All data supporting the findings of this study are available within the paper. Mass spectrometry data have been deposited in ProteomeXchange with the primary accession code PXD053136 and PXD053139. This paper does not report any original code. All other data supporting the findings of this study are available from the corresponding author on reasonable request. Source data are provided with this paper.

